# One-Day Construction Of Multiplex Arrays to Harness Natural CRISPR Systems

**DOI:** 10.1101/2020.03.06.981316

**Authors:** Robert M. Cooper, Jeff Hasty

**Affiliations:** BioCircuits Institute, University of California, San Diego, La Jolla, CA; San Diego Center for Systems Biology, La Jolla, CA; Molecular Biology Section, Division of Biological Sciences, University of California, San Diego, La Jolla, CA; Department of Bioengineering, University of California, San Diego, La Jolla, CA

## Abstract

CRISPR-Cas systems are prokaryotic immune systems that have proliferated widely not only in bacteria and archaea, but also much more recently, in human biological research and applications. Much work to date has utilized synthetic sgRNAs along with the CRISPR nuclease Cas9, but the discovery of array-processing nucleases now allows the use of more compact, natural CRISPR arrays in heterologous hosts, in addition to organisms with endogenous systems. Unfortunately, the construction of multiplex natural CRISPR arrays remains technically challenging, expensive, and/or time-consuming. This limitation hampers research involving natural CRISPR arrays in both native and heterologous hosts. To address this problem, we present a method to assemble CRISPR arrays that is simple, rapid, affordable, and highly scalable – we assembled 9-spacer arrays with one day’s worth of work. We used this method to harness the endogenous CRISPR system of the highly competent bacterium *Acinetobacter baylyi*, showing that while single spacers are not always completely effective at blocking DNA acquisition through natural competence, multiplex natural CRISPR arrays enable both nearly complete DNA exclusion and genome editing, including with multiple targets for both. In addition to demonstrating a CRISPR array assembly method that will benefit a variety of applications, we also find a potential bet-hedging strategy for balancing CRISPR defense vs. DNA acquisition in naturally competent *A. baylyi*.

---

CRISPR (clustered regularly interspaced short palindromic repeats)-Cas systems are adaptive immunity mechanisms that protect bacteria and archaea against invading nucleic acids, generally by detecting and cutting defined target sequences^1^. CRISPR systems include Cas (CRISPR-associated) proteins, as well as their eponymous arrays of short direct repeats that alternate with similarly short DNA spacers. The spacer array is transcribed into a long pre-crRNA, which is then processed into individual crRNAs (CRISPR RNAs), each composed of a single spacer that is complementary to a particular nucleic acid target, and often a hairpin handle derived from a repeat. These crRNAs bind Cas effector proteins, such as Cas9, or protein complexes, such as CASCADE. Once bound, they guide the effector to complementary DNA or RNA, depending on the system, which the effectors often cleave.

In short order, many labs have adapted CRISPR-mediated DNA cleavage for applications ranging from precise genome engineering to genetic circuits^2^ to targeted bacterial strain removal^3-6^. Self-spreading CRISPR constructs have also been used to quickly generate homozygous diploid knock-outs (the mutagenic chain reaction)^7^, and preliminary work suggests they could re-engineer entire populations through biased inheritance; i.e., gene drives or active genetics^8-13^.

Spacer multiplexing is beneficial for nearly all of these applications^14^. Targeting multiple sites on the same gene improves both mutagenesis and gene regulation^2^, cleaving multiple target sites prevents emergence of resistant alleles^15^, and multiple genes can be edited simultaneously. While natural CRISPR arrays are inherently multiplex – some including hundreds of spacers – multiplexing in synthetic biology applications has been comparatively limited. One reason is that constructing synthetic multiplex CRISPR arrays is technically challenging due to their extensive repetition. Addressing this difficulty, several strategies have been developed to assemble tandem arrays of synthetic sgRNA (single guide RNA) transcriptional units, but these were limited in array size or required time-consuming, sequential cloning for each additional spacer^16-19^. Recently, others have shown that single-promoter sgRNA arrays can be assembled using tRNAs to direct processing and release of individual sgRNAs^20,21^.

Natural CRISPR arrays were largely abandoned in favor of such synthetic sgRNAs because this system only requires a single effector protein for use in non-native hosts, namely Cas9. However, the more recent discovery that other single-protein CRISPR effectors, including Cas12a (Cpf1) and Cas13a (C2c2), can process natural arrays without tracRNA means that natural, multiplex CRISPR arrays can be used in non-native hosts as easily as sgRNAs^22-24^. In comparison to artificial sgRNA arrays, natural CRISPR arrays have several advantages for multiplexing. Natural arrays are much more compact, making them easier to package and deliver. Natural arrays also have a particular advantage for applications in prokaryotes, many of which already have their own endogenous CRISPR systems that can be retargeted using synthetic spacers^1^. One attractive application could be to limit horizontal gene transfer, a major contributor to multi-drug resistance and pathogenicity^25^.

Unfortunately, the signature palindromic repeats significantly complicate assembly of natural CRISPR arrays. This problem is particularly important because spacer design rules are not completely accurate even for the best studied Cas nucleases, so developing good arrays can require building and testing multiple designs^26^. Recent approaches for assembling multiplex natural arrays have been limited to just a few spacers^4,27^, imposed sequence constraints^28^, or required sequential, time-consuming cloning steps for each additional spacer^5,28,29^. Multiplex arrays can be assembled using very long single-stranded oligos (180 nt^23^), but these become significantly more expensive and unreliable as their length surpasses 60 nt. Another option is double-stranded DNA synthesis^5,24^, but this can also be unreliable or require slower, more expensive cloned gene services. Such double-stranded DNA synthesis often takes longer or fails for sequences containing repetition and/or secondary structure^24,30^, both of which are defining features of CRISPR arrays. Primed adaptation can generate multiplex arrays using the endogenous adaptation mechanism, but the results are stochastic, not designed^31,32^. A recent one-pot method enables rapid assembly of nearly-natural CRISPR arrays, but this still requires trimming the 3’ ends of spacers^33^. This makes the method incompatible with systems that do not trim their spacers and thus require sequence complementarity throughout, including the most prevalent Type I systems^1^. Array assembly therefore remains a key challenge in the field^14^.

As a testbed for multiplex harnessing of an endogenous CRISPR system, we used the highly naturally competent bacterium *Acinetobacter baylyi*^34^, which is a largely non-pathogenic, but phenotypically similar, relative of the highly drug-resistant and clinically urgent pathogen *A. baumannii*^35-37^. *A. baylyi* is an ideal platform for developing endogenous CRISPR applications, because it combines extensive natural competence, which promotes horizontal gene transfer, with a CRISPR system that prevents it. We first demonstrated that *A. baylyi* has a functional Type I-F CRISPR system, but that it is not fully effective against gene acquisition by natural competence when using single spacer arrays. To better harness the endogenous system, we developed a method to construct natural CRISPR arrays that is rapid (one day), scalable (we had no difficulty reaching lengths of 9 spacers), affordable, and results in completely natural arrays with no sequence modifications, constraints, or considerations.

## Results

### *A. baylyi* contains a functional Type I-F CRISPR system

The *A. baylyi* genome contains a computationally identified Type I-F CRISPR system (Figure 1A)^37^, but its function has not been tested experimentally. Therefore, we first determined whether the endogenous CRISPR system can block horizontal gene transfer via natural competence. To test the system, we inserted single-spacer arrays **t**argeting a kanamycin resistance gene into a previously used neutral locus in the genome^38^. We tested four different spacers from both the top (T) and bottom (B) strands, each using the 5’-CC-protospacer-3’ protospacer-adjacent motif (PAM, 5’-anti-protospacer-GG-3’ on the complementary, targeted strand) previously shown to work in the Type I-F systems of *E. coli*^39^, *Pectobacterium astrosepticum*^40^, and *Pseudomonas aeruginosa*^41^. When naturally competent cells carrying these single arrays were incubated with a self-replicating plasmid (pBAV-K1)^42^, there were still many kanamycin-resistant transformants, and only the T2 spacer reduced the transformation efficiency relative to a random spacer (Figure 1B, note the log scale). When they were challenged using a genomically integrating linear DNA construct (Vgr4-K1), again the T2 spacer worked well, now decreasing acquisition of kanamycin resistance by 1000-fold relative to a random spacer, but the others were less effective (Figure 1C). Escape clones did have somewhat smaller colony sizes, suggesting partial tolerance for ongoing self-targeting. All strains remained competent for Vgr4-K2, which contains a second kanamycin resistance gene with minimal homology to the first (Figure 1D).

**Figure 1:**
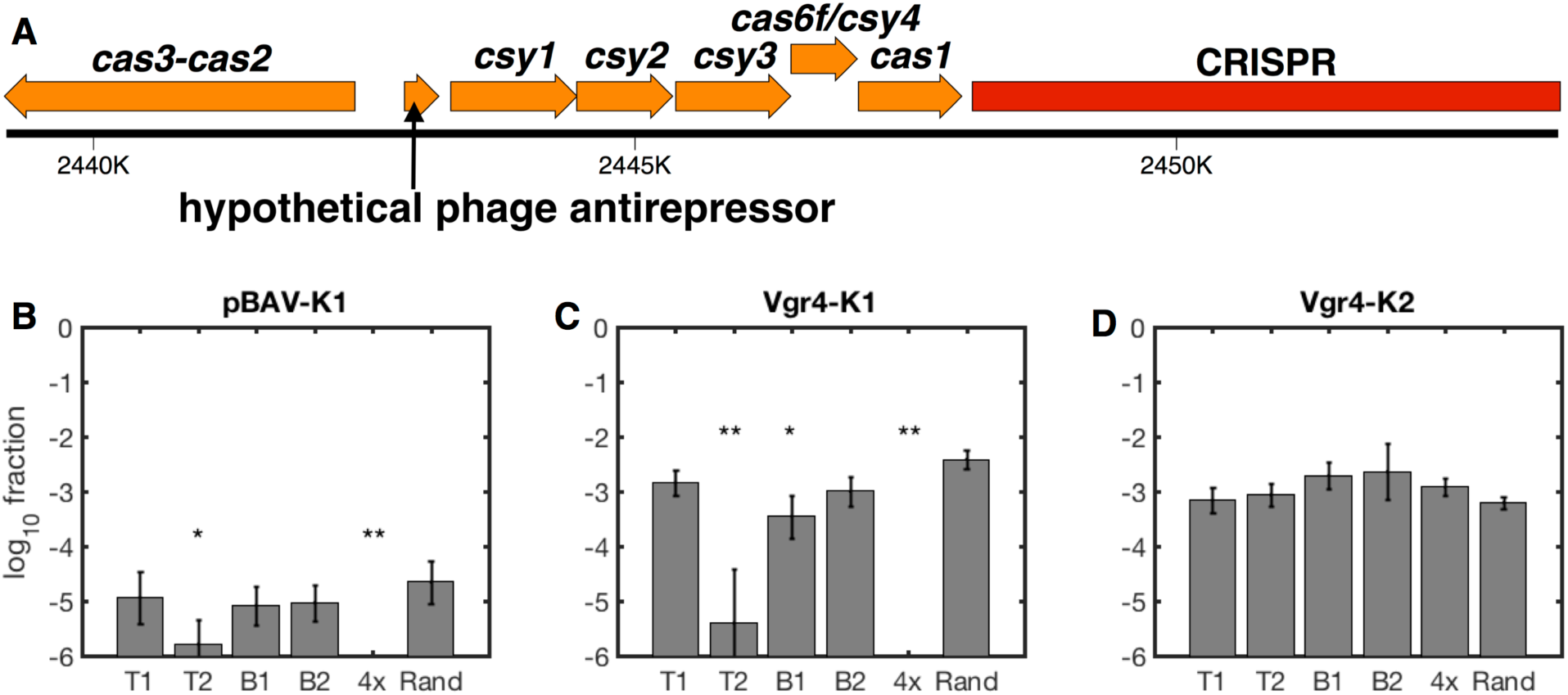
Synthetic *A. baylyi* CRISPR arrays blocking gene acquisition via natural competence. (A) The endogenous, Type I-F CRISPR locus in *A. baylyi*. (B-D) Cells containing individual spacer arrays (T1, T2, B1, or B2), a 4-spacer multiplex array including all individual spacers, or a random spacer were naturally transformed with the self-replicating plasmid pBAV-K1 (B), the integrating linear DNA Vgr4-K1 (C), or the non-targeted, integrating linear DNA Vgr4-K2 (D). Note the fraction of cells acquiring kanamycin resistance is shown on a log scale. Data includes 2 experimental replicates, each with 3 measurement replicates, error bars indicate propagated standard deviations (see Methods), and limits of detection were roughly 10^−6^. Statistical comparison to the random spacer was performed using multiple comparison analysis (Methods): *=*p*<.01, **=*p*<10^−6^.

### Construction of Multiplex CRISPR Arrays

To increase the efficacy of the endogenous *A. baylyi* CRISPR system against incoming DNA, we next turned to multiplex arrays, which have been reported to increase CRISPR efficacy in a variety of contexts. However, constructing natural, multiplex Type I CRISPR arrays remains challenging for the reasons described above. Therefore, we developed a new method to assemble multiplex, completely natural arrays.

Our method is based on annealing and ligating single-stranded DNA oligos (Figure 2). The key insight is that despite extensive repetition, the correct order can be ensured by avoiding annealing or ligation within repeats. To achieve this, we design 60 nt top oligos that each include a single 28 nt repeat in their center and extend halfway (16 nt) into the spacer or flanking sequence on either side. These top oligos are joined together by annealing to 40 nt bottom bridge oligos, consisting of the reverse complement of each 32 nt spacer plus 4 nt of repeat on either side. The intentional gaps on the bottom strand avoid oligo annealing within repeats, and they are filled in later by PCR. We tested multiple conditions in optimizing our assembly protocol (Figure S2), and our final optimized protocol is as follows (see Methods for more detail):

**Figure 2:**
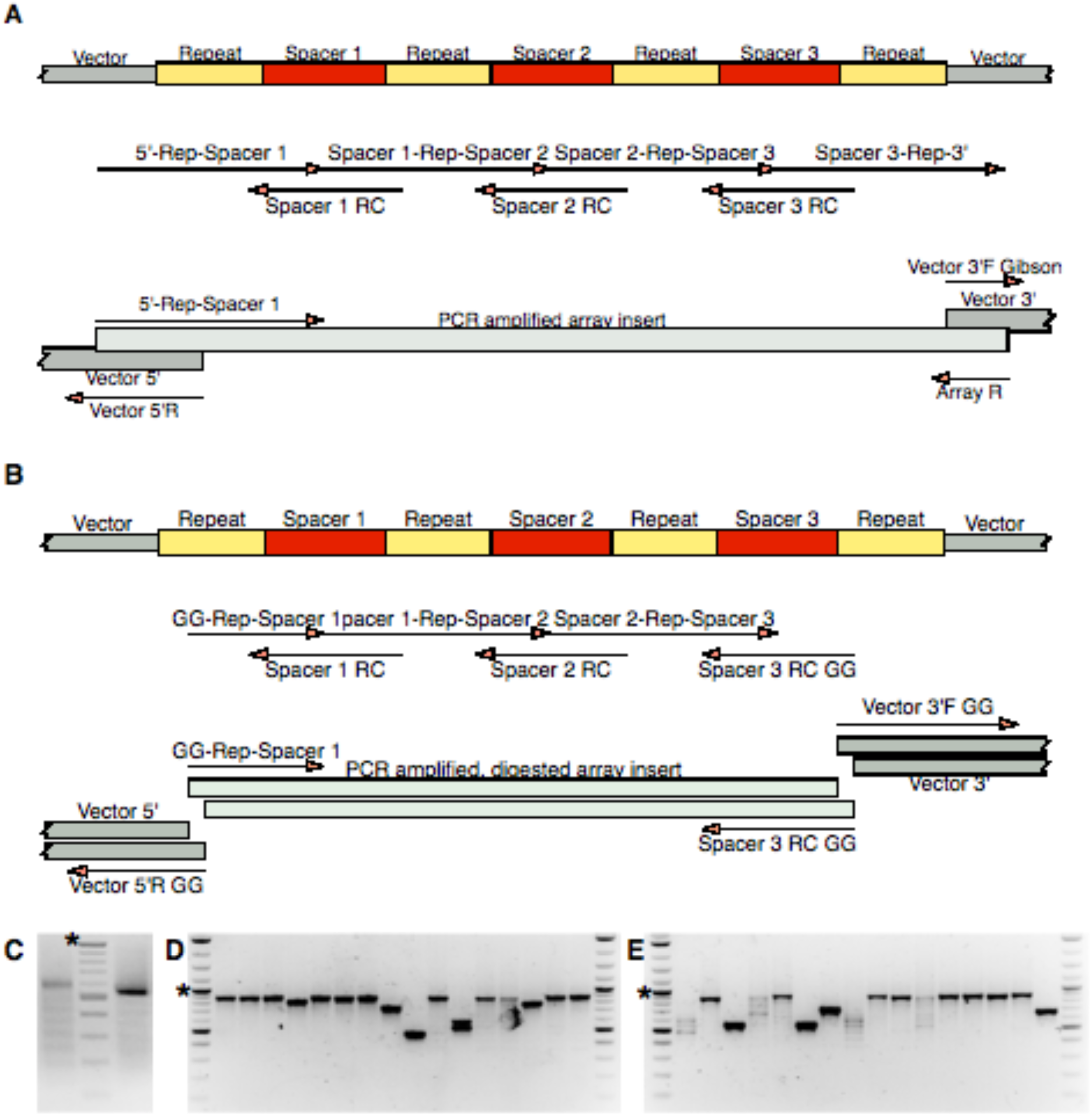
Strategy for Assembling Multiplex, Natural CRISPR Arrays. Assembly strategy for a sample 3-spacer CRISPR array to be inserted into a vector using Gibson assembly or fusion PCR (A), or Golden Gate assembly (B). Each strategy shows the desired end product, the top and bottom oligos used for array annealing and ligation, and the PCR amplicons for insertion into a vector. Single-stranded primers (oligos) are shown as arrows pointing 5’ to 3’. Note that primers used for Golden Gate assembly (denoted “GG”) have an additional Golden Gate tail appended to their 5’ ends (see Methods). See Supplementary Figure S1 for more detailed versions that include DNA sequence. C) PCR amplified 9-spacer arrays using the Gibson (left) and Golden Gate (right) strategies. Colony PCR screening of *E. coli* clones for 9-spacer arrays inserted using Golden Gate (D) and Gibson (E) strategies, where the correct length is 914 bp. The ladder on all gels has 100 bp increments, with the 1 kb band marked by an asterisk.

1. <u>Phosphorylation</u>: Mix 2 to 4 ul of each top oligo from 100 µM stock solutions (Supplementary Figure S1A), and phosphorylate them using T4 polynucleotide kinase (PNK) and 1x T4 DNA ligase buffer at 37°C for 15-60 minutes. This step can be skipped if ordering 5’ phosphorylated oligos. Phosphorylating the top oligos separately increases PNK activity, which is optimal on single-stranded DNA.
2. <u>Annealing</u>: Mix 1 part top oligos with 2-3 parts bottom oligos by molarity (Supplementary Figure S1B), and perform a slow annealing starting from 90°C. We used a thermocycler programmed to decrease to 37°C by 0.1°C/sec, but allowing a hot water bath to gradually cool should work as well.
3. <u>Ligation</u>: Add T4 DNA ligase and additional ligase buffer, and incubate at 37°C for 30 minutes.
4. <u>Clean up</u>: Column purify the ligated array using a standard DNA purification column to remove unincorporated oligos.
5. <u>Amplification</u>: PCR amplify the array using primers appropriate for your cloning strategy of choice, *e.g.*, Gibson or Golden Gate assembly, using as high an annealing temperature as the primers will allow (Supplementary Figure S2C-E).
6. <u>OPTIONAL: Gel Purification</u>: Run the raw ligation or amplified PCR product on an agarose gel, excise the correct band, and purify the DNA using a gel extraction kit. This step is optional for shorter arrays, but it can substantially increase accuracy for longer arrays.
7. <u>Insert into vector</u>: Insert the array into a vector using a method of your choice; *e.g.*, Golden Gate, Gibson assembly, or fusion PCR.
8. <u>Transform</u>: Transform the final construct into *E. coli* (for circular plasmids), or directly into *A. baylyi* (for linear constructs with genomic homology for recombination), spread on selective agar plates, and incubate overnight.
9. (<u>Next day) Screen</u>: On the following day, pick several colonies and PCR across the array to screen for assemblies of the correct length (Supplementary Figure S2D,E).

The assembly steps can be completed in one day, and the resulting colonies can be screened the following day by PCR across the CRISPR array. This basic array assembly technique is compatible with multiple cloning strategies for insertion into a final vector. In developing our protocol, we successfully inserted the arrays into circular plasmids using both Gibson (Fig. 2A and S1A) and Golden Gate (Fig. 2B and S1B) cloning strategies, as well as into linear DNA fragments that we amplified via PCR.

Using our optimized protocol, we were able to quickly and accurately assemble a 9-spacer array (Figure 2C), using either Gibson or Golden Gate strategies to insert the array into the plasmid. For Golden Gate insertion, 11 of 16 picked colonies had the correct length array (Figure 2D), and for Gibson insertion, 8 of 16 picked colonies had the correct length (Figure 2E). Sanger sequencing confirmed that all arrays with the correct length were assembled in the correct order. 7 of the Golden Gate and 2 of the Gibson clones were completely correct, and the remainder had various indels or substitutions. Only one of the errors was at a junction between oligos, suggesting most may have occurred during oligo synthesis.

### Multiplex Natural Arrays Enhance CRISPR Efficacy In Natural Competence

To see if multiplex CRIPSR arrays more effectively interfere with natural competence in *A. baylyi*, we combined the 4 spacers targeting the kanamycin resistance gene into a single, 4-spacer natural array and inserted it into the *A. baylyi* genome. This 4xKan1 array was highly effective against both the self-replicating plasmid pBAV-K1 and the genomically integrating construct Vgr4-K1 (Figure 1B,C, Figure 3A). As for single-spacer arrays, the 4-spacer array was ineffective against a second, control kanamycin resistance gene with no homology to the targeted gene (Figure 1D, Figure 3B). The 4-spacer array allowed no escape transformants with the replicating plasmid, but we did obtain 2 escapes with the integrating construct. In one of these escapes, the inserted 4-spacer CRISPR array had been disrupted by the active insertion sequence IS1236^43^. The other escape appeared to have a larger genomic deletion encompassing the array, as it had lost the spectinomycin resistance marker used to select for insertion of the array, and the entire region failed to amplify by PCR.

**Figure 3:**
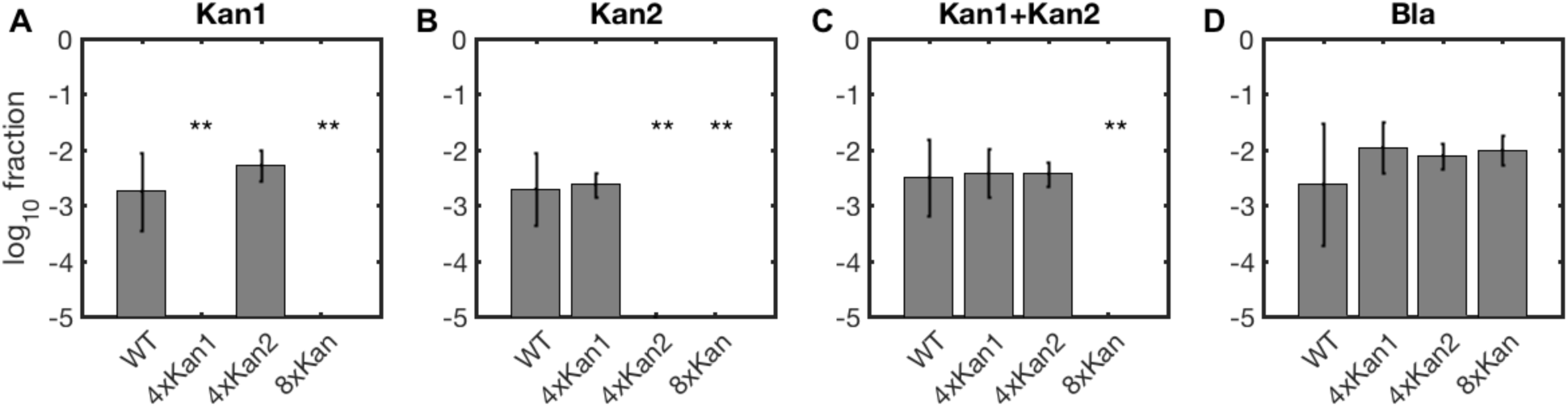
Synthetic, multiplex *A. baylyi* CRISPR arrays block acquisition of multiple genes. Cells containing no exogenous CRISPR arrays (WT), 4-spacer arrays targeting *kan1* and *kan2*, and an 8-spacer array targeting both *kan* genes (x-axis tick labels) were incubated with linear, genomically integrating DNA. Donor DNA constructs included Vgr4-Kan1 (A), Vgr4-Kan2 (B), both *kan* constructs (C), or a non-targeted beta-lactamase gene (D). Data includes 2 experimental replicates, each with 3 measurement replicates, error bars indicate propagated standard deviations (see Methods), and limits of detection were roughly 10^−6^. **=*p*<10^−7^.

Next, we expanded our array to defend against both kanamycin resistance genes simultaneously, using an 8-spacer array. As a preliminary step, we constructed a 4-spacer array targeting the second kanamycin gene, added genomic homology arms via fusion PCR, and cloned the linear product into *A. baylyi*. Then we assembled an 8-spacer array targeting both kanamycin resistance genes. We easily assembled this 8-spacer array in a one-pot reaction, but we also assembled it from the individual 4-spacer arrays to demonstrate modular array construction. For the modular approach, we PCR amplified the cloned *4xKan2* array using a leftmost top primer that began with the first 16 bp of the final spacer in the *4xKan1* array rather than with the 5’ region of the vector, and then performed a fusion PCR of the 3 pieces *Vector 5’-4xKan1, 4xKan2*, and *Vector 3’*.

In contrast to single spacers (Figure 1), each 4-spacer array effectively blocked acquisition of its respective kanamycin resistance gene (Figure 3A,B), and only the 8-spacer array prevented acquisition of kanamycin resistance when both genes were present (Figure 3C). All arrays allowed acquisition of a non-homologous beta-lactamase gene (Figure 3D). The modular construction shows that even if there is a size limit to this method, very large arrays can still be assembled in very few steps.

### Markerless Genome Editing Using Endogenous CRISPR Systems

CRISPR has been used for genome editing in many contexts, and we wanted to confirm that our natural arrays would enable editing of the *A. baylyi* genome as well. To do this, we constructed a 3-spacer array targeting the *bap* gene (ACIAD2866), which has been implicated in biofilm formation in *Acinetobacter*^44,45^, and thus may be at least partially responsible for intractable clogging when using *A. baylyi* in microfluidics^46^. We inserted the *3xBAP* array into both pBAV1spec for cloning into *E. coli*, as well as into a linear construct with roughly 1 kb genomic homologies on either side for direct insertion into the *A. baylyi* genome. The pBAV1spec assembly transformed into *E. coli* was the correct length in 8 of 8 tested clones (Figure 4A, left half). We sequenced four of them, of which all had the correct spacer order, although one was missing two base-pairs. When we co-transformed this pBAV1spec-CRISPR_3xBAP_ into *A. baylyi* along with a markerless *bap* deletion donor DNA (linear dsDNA with ∼1 kb homology arms on either side), both of two tested clones had the correct deletion (Figure 4B). Interestingly, we found that *bap* in our strain of *A. baylyi* ADP1 (ATCC 33305) was approximately 3 kb larger than in the published genome. This may have been due to a sequence assembly error or genomic instability, either of which could result from the many tandem repeats found in *bap* genes^44,45^, but we did not pursue the discrepancy.

**Figure 4:**
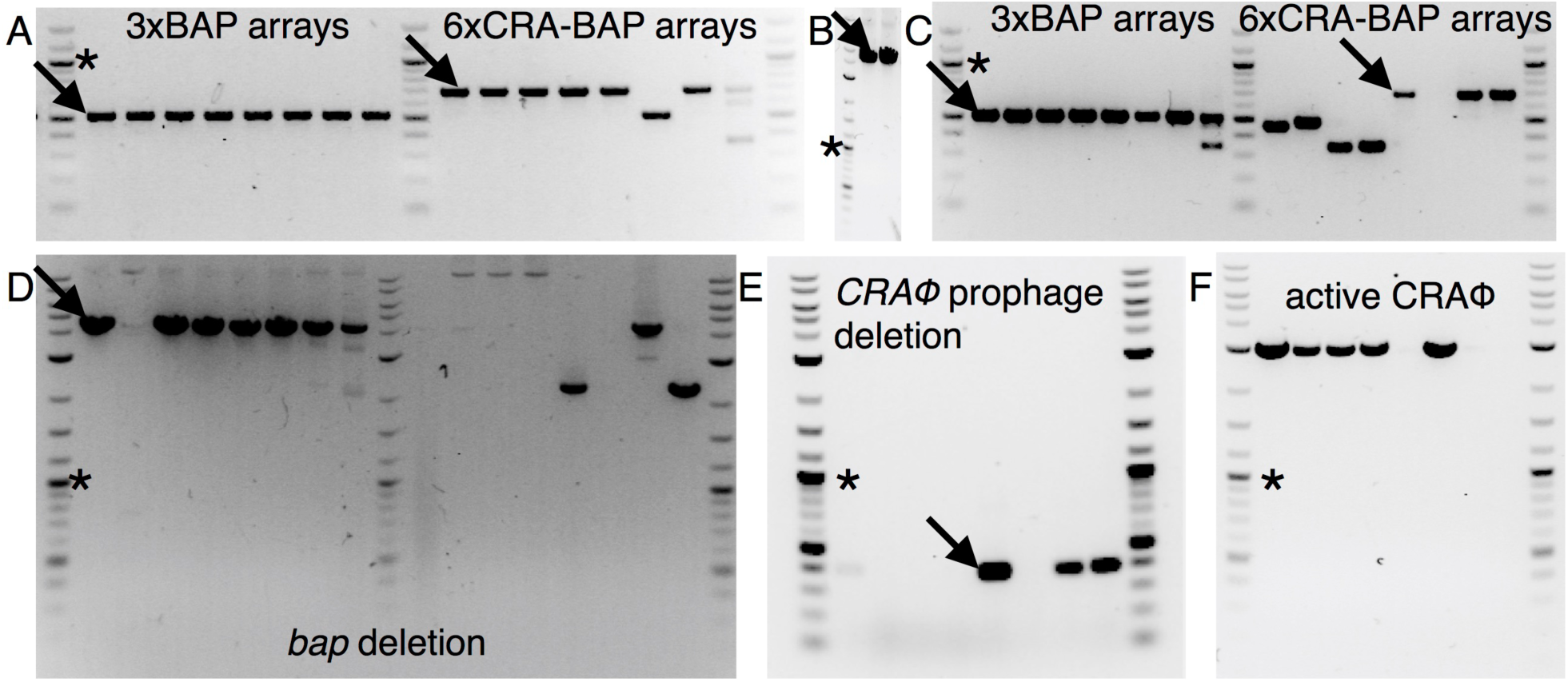
Multiplex genome editing using endogenous CRISPR systems. Arrows indicate the expected bands for correct genomic edits, and asterisks indicate the 1 kb band of the ladder (not counted in lane numbering). A) PCR screening of of 3xBAP (lanes 1-8) and 6xCRA-BAP (lanes 9-16) arrays in pBAV, cloned into *E. coli*. B) PCR screening of 2 markerless *bap* deletions in *A. baylyi* using pBAV-CRISPR_3xBAP_. C-F) PCR screening of markerless *bap* and double CRAΦ, *bap* deletions in *A. baylyi* using non clonal, linear PCR products from array assembly. C) Multiplex 3xBAP (lanes 1-8) and 6xCRA-BAP (lanes 9-16) arrays. D) *bap* deletion screening for the same clones as in C. The deletion and wild type amplicons are roughly 4.5 and 12 kb, respectively. E) CRAΦ deletion screening for the clones in lanes 9-16 of C,D. Product was only expected for CRAΦ deletion. F) As in E, but circular CRAΦ phage screening. The 3 kb product was only expected if CRAΦ was present in its excised, circular episome form.

When using a linear construct to deliver the *3xBAP* array into *A. baylyi*, we obtained far more clones than when using pBAV1spec (on the order of 1000 vs 36), which is expected because homologous recombination is more efficient than plasmid re-circularization in *A. baylyi* natural competence^47^. Of 8 tested clones, 7 had the correct size array (Figure 4C, left half) and 7 had the correct BAP deletion (Figure 4D, left half), even despite the CRISPR array not having first been clonally verified.

Next, we attempted to delete two genes at once, creating a 6x array targeting both *bap* and the CRAΦ prophage, which binds the competence machinery when activated, complicating horizontal gene transfer experiments^48^. The pBAV1spec-CRISPR_6xCRA-BAP_ construct had the correct array length in 6 of 8 *E. coli* clones (Figure 4A, right half), but we were unable to use it to obtain a double genomic deletion in *A. baylyi*, likely due to the relative inefficiency of circular plasmids in natural transformation.

To increase our triple transformation efficiency, we used the genomically integrating, linear *6xCRA-BAP* construct along with CRAΦ and *bap* deletion donor DNAs. Of 8 tested clones, 3 had the correct array length (Figure 4C, right half). All 3 of those had both the desired genomic CRAΦ deletion (Figure 4E) and eliminated the excised, circular CRAΦ episome (Figure 4F). All three clones also had mutations in *bap*, although two of them had larger deletions (Figure 4D, right half), leaving one clone with both precise deletions. One of the larger *bap* deletions extended to the end of a nearby copy of the insertion sequence IS1236^43^, and the other had a more complex rearrangement that appeared to involve an inversion of part of the genome. IS1236 is not present next to *bap* in the official genome sequence, but it was already there in our parental strain before the double deletion attempt. This is not completely unexpected, since IS1236 is known to be highly active in *A. baylyi*^49^. If the correct editing rate were more important than speed, one could likely increase the percentage of clones with the correct edits by first clonally verifying the linear CRISPR array construct.

### Construction of Cas12a Arrays

Finally, we demonstrated that the method described here is generalizable to other natural CRISPR arrays, which use different repeat sequences and spacer lengths. For this demonstration, we chose Cas12a/Cpf1 arrays, which are processed by their respective single effector nuclease^50^. The Cas12a CRISPR array unit for *Franciscella novicida* U112 is slightly longer than the *A. baylyi* array unit, with 36 bp repeats and 26-32 bp spacers^51^. Nevertheless, we easily assembled a 4-spacer array with a full 68 bp unit length, targeting a beta lactamase gene (Supplementary Figure S3A). All screened clones (8 of 8) had the full-length array in the correct order (Supplementary Figure S3B) of which 2 were correct with no gaps.

## Discussion

The method presented here solves the challenge of rapid, affordable, and scalable construction of completely natural multiplex CRISPR arrays, with no sequence modifications and only minimal constraints. This should be highly beneficial for multiple applications in a variety of organisms, from basic research to applied tools. For applications using heterologous, array-processing Cas nucleases such as Cas12a, facile construction of multiplex natural arrays will help with gene regulation^22^, genome engineering^23,24^, and even population engineering^15^.

We designed this assembly method to include 3 key features that improve its accuracy and efficiency: unique ligation junctions, long annealing regions, and limited oligo length. In the first feature, the only ligation junctions are within the unique spacers on the top strand, which helps to ensure assembly in the correct order. We purposely left gaps in the repeat regions on the bottom strand to avoid ligation junctions within repeats. We tested including an oligo covering the remaining 20 nt of the repeats to fill in the gaps on the bottom strand (repeat_RC), but this resulted in a smear of larger than expected ligation products, indicating increased ligation at incorrect junctions (Fig. S2A). Furthermore, while developing this protocol we sequenced several correct-sized clones that had incorrect spacer order, but only when including the repeat_RC oligo.

The second feature is long (20 nt) annealing regions that allow more rapid and specific annealing and ligation than the usual 4 bp Golden Gate overlaps, particularly at the 37°C where T4 DNA ligase has optimal activity. The long annealing regions also allow the user to choose spacers without constraints imposed by the requirement for junction orthogonality, since such long sequences should be highly specific. This allows for very easy, plug-and-play oligo design. Third, the longest oligos must only be the unit length of the CRISPR array, which for *A. baylyi* is 60 nt. Oligos of this length are relatively reliable, affordable, and rapidly delivered from most DNA synthesis vendors.

A final advantage lies in cost-saving oligo reusability. Unlike *ad-hoc* construction strategies, this method places the ligation junctions in the same location for every spacer-repeat unit, meaning that many oligos can be reused for alternate array designs without checking for compatibility. For example, our *4xKan1* and *4xKan2* arrays were easily joined with just one additional oligo. This modular assembly demonstrates that verified sub-arrays can easily be joined with just one additional day of work.

It should be noted that this method does have a few limitations. First, while it can generate arrays of defined spacer order, it cannot generate randomized array libraries. Second, arrays with multiple copies of the same or highly homologous spacers could be challenging to assemble. Third, spacers that have strong secondary structures or that are complementary to either other spacers or the repeat could pose a challenge.

The PCR amplification step following ligation both enriches the correct size product and produces a double-stranded construct with no gaps. A fully double-stranded insert is particularly important for Gibson Assembly-based insertion into the vector because of the required exonuclease, but we also found it to be important for Golden Gate insertion. Without PCR amplification, Golden Gate insertion of a 6x array yielded clones containing a range of incorrectly sized inserts (compare Supplementary Figures S2D and S2E). Interestingly, these incorrect arrays almost always contained spacers that were in the correct order, but truncated at the 5’ end. We suspect the 5’-specific truncation may involve a gap repair process within the *E. coli* host that may be mediated by repeats and directionally biased by plasmid replication.

In prokaryotes with endogenous CRISPR systems, this method will improve the study and understanding of the ecological importance of CRISPR in its natural context, including the antagonistic interplay between CRISPR and horizontal gene transfer (HGT)^25,52-54^. This seemingly contradictory pair of abilities has raised evolutionary questions about tradeoffs between the acquisition of new traits via HGT, versus CRISPR-mediated exclusion of foreign DNA^55,56^. This interaction is important for microbial evolutionary theory, but when the transferring genes confer antibiotic resistance or pathogenicity, it also directly impacts human health^57^. Here, we demonstrated that in the highly competent *A. baylyi*, the CRISPR-HGT interaction is not straightforward. While multiplex arrays effectively blocked exogenous DNA uptake, weaker single spacers reduced, but did not eliminate, HGT. This suggests that for *A. baylyi*, one solution to the CRISPR-HGT conundrum is to hedge their bets. Single spacers provide some protection against incoming targeted DNA, but particularly for weaker spacers or when multiple spacers compete for limited CASCADE complexes^58^, some targeted DNA can still be acquired. When the tolerance is only partial, the targeted protospacer (or the CRISPR machinery) will eventually mutate to eliminate genomic self-targeting and alleviate growth costs, allowing ongoing exploration of the genetic diversity in the environment.

## Methods

### Array construction

We designed spacers to match target sequences preceded by CC on the non-targeted strand using a computational tool to ensure they were maximally orthogonal to the rest of *A. baylyi* genome^59^. Briefly, the algorithm searches for all possible spacers in the target sequence that have the appropriate PAM, and then scans them against the host genome to find the most similar sequence, giving greater weight to bases in the PAM-proximal seed sequence. The best match (highest score) against the host genome is assigned as the score for that spacer. We chose spacers from among the lowest scoring (most genome-orthogonal) sequences to cover the entire target and include both DNA strands. For a random spacer, we found the lowest scoring sequence among a computer-generated, random pool. Oligos were designed according to the diagrams in Figure 2 and Supplementary Figure S1, and their sequences are given in Supplementary Table 1. Spacer sequences are shown in Supplementary Table 2. We ordered standard quality, desalted oligos normalized to 100 uM in TE buffer from ValueGene, Eton Bio, and Integrated DNA Technologies. All enzymes and buffers were from New England Biolabs. To construct arrays, we used the following optimized procedure:

1. Phosphorylate oligos by mixing 1-2 ul of each top-strand oligo along with 1x T4 ligase buffer and 1 ul T4 polynucleotide kinase (NEB). Polynucleotide kinase buffer will not work without supplementary ATP. Incubate at 37 degrees for 30-60 minutes.
2. Anneal oligos by mixing 1 part phosphorylated top oligos with 2 to 3 parts bottom oligos, heating to 85°C, and slowly cooling back to 37°C at 0.1°C per second in a thermocycler.
3. Ligate by adding 1 ul T4 DNA ligase and another 1x ligase buffer. Incubate at 37°C for another 30-60 minutes.
4. Remove unligated oligos using a PCR purification column (Lamda Biotech).
5. PCR amplify the ligation product using primers as shown in Figures 2 and Supplementary 1. We used Q5 DNA polymerase and the manufacturer’s recommended protocol, annealing at 72°C, extending for 20 seconds, and running for 20 cycles. A high annealing temperature is critical to recover the correct product; primers can be checked at http://tmcalculator.neb.com/.
6. Purify the PCR product either directly or after excising the correct band from a gel, using a column-based PCR or gel purification kit (Qiagen).
7. Insert the array into a vector. For Gibson assembly, we mixed 2 ul total DNA (with equimolar parts) with 2 ul of 2x master mix and incubated at 50°C for one hour. For Golden Gate assembly, we mixed 4 ul total DNA (with equimolar parts), 0.5 ul T4 DNA ligase buffer, 0.25 ul T4 DNA ligase, and 0.25 ul BsaI, and incubated for 30-50 cycles of 1 minute each at 37°C and 24°C, followed by 10 minutes at 50°C. Vectors were prepared by PCR using primers as shown in Figures 1 and S2, and gel extracted. Whenever the vector PCR was derived from a plasmid, we used the primers Vector 3’F and Vector 5’R and treated the product with DpnI. For linear constructs used in direct transformation into *A. baylyi*, the vector consisted of approximately 1 kb homology arms on either side of the array. In these cases, we either directly mixed the 3 pieces (5’ arm, array, and 3’ arm) in a full-length PCR reaction, or first pre-joined the 3 pieces via either Gibson or Golden Gate assembly, and then PCR amplified and gel extracted the full construct.

For modular assembly of the *8xKan* array, we first assembled both *4xKan1* and *4xKan2* arrays and inserted them into the genomic integration vector as above. Next, we PCR amplified the 5’ part of the *4xKan1* construct through the array using the primers pp_5’F and Kan1_B2_RC, as well as the *4xKan2* construct using using the primers Kan1_B2-R-Kan2_T1 and Array_R. Then we performed a 3-piece PCR with primers Vector_5’F and Vector_3’R to fuse (i) Vector 5’-4xKan1, (ii) 4xKan2, and (iii) the vector 3’ piece (amplified using primers Vector_3’F and pp_3’R).

To assemble FnCas12a arrays, we followed the same procedure described above, using the Golden Gate insertion strategy.

### Cell culture, transformations, and screening

We grew all cells in LB media at 30 or 37°C. *A. baylyi* strain ADP1 was obtained from ATCC (stock #33305) and for *E. coli* we used a lab strain of MG1655. The *kan1* gene was aminoglycoside O-phosphotransferase APH(3’)-IIIa, and the *kan2* gene was aminoglycoside O-phosphotransferase APH(3’)-IIa. These two genes have no significant similarity as determined by BLAST alignment. For transformation of *A. baylyi* via natural competence, we washed overnight cultures, resuspended in fresh LB, and incubated 50 ul of cells plus DNA at 37°C for 2 to 4 hours. All data plotted in the same figure used the same concentration of donor DNA, generally 0.2-1 ng/ul. To quantify the fraction of transformed cells, we performed five 10-fold serial dilutions and spotted 3 measurement replicates of 2 ul each at each dilution level onto 2% agar LB plates containing the appropriate (or no) antibiotic selection (20 ug/ml of kanamycin and/or spectinomycin). Each experiment was repeated on two separate days. Lower agar concentrations did not work well for colony counting, because the motile cells began to spread and colonies became less well-defined. Only colonies visible after 20 hours at 30°C for 20 hours were counted.

We inserted CRISPR arrays into a neutral genomic region that has been used previously, replacing genomic coordinates 2,159,575-2,161,720, covering ACIAD2187, ACIAD2186 and part of ACIAD2185^38^. The integration site for CRISPR-targeted kanamycin resistance genes was another region that we have found to be neutral in our lab conditions, ACIAD3427. The upstream homology arm covered coordinates 3,341,420-3,342,480, and the downstream homology arm covered 3,342,641-3,343,720. Our replicating plasmid was the broad host pBAV1k^42^, which we modified to spectinomycin resistance when using it to carry CRISPR arrays. In our arrays, we included the 80 bp upstream of the endogenous CRISPR array to include any leader sequences or regulatory elements. For markerless genomic deletions, we constructed linear donor DNA by PCR fusing approximately 1 kb regions upstream and downstream of the targeted gene.

For PCR screening of clonal CRISPR arrays in *E. coli*, we picked individual colonies into 50 ul of water, and used 1 ul directly in a PCR reaction. For *A. baylyi*, we found we did not obtain clean results unless we first used a genomic miniprep kit to purify DNA (Promega Wizard). We inverted the colors for all agarose gels to assist visualization.

### Statistical analysis

To calculate error bars for ratios on logarithmic plots, we used error propagation as described previously^46^. For each experimental replicate (each with 3 measurement replicates; i.e., 2 *ul* spots), we took the log base 10 of each data point, found the standard deviations for both transformed and total cell count measurement replicates (*σ*_1_ and *σ*_2_), and calculated the standard deviation of the ratio as 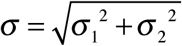. To find the total variance across experimental replicates from different days, we used the error propagation formula 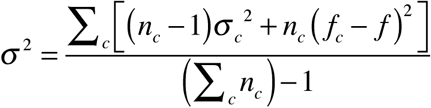, where the subscript *c* denotes experimental replicates, *f* is the fraction transformed, and *n*_*c*_ is the number of measurement replicates for each experiment (here, 3 spotting replicates). Performing calculations on a logarithmic scale creates a problem when some, but not all, measurement replicates are below the limit of detection, because zeros create infinities. In these cases, we set the zeros to half the limit of detection as a conservative estimate for the purposes of plotting, since excluding them would artificially increase the average for that experiment.

We performed significance tests as described previously^46^. In Figures 1 and 3, we performed multiple comparison tests using the Matlab function multcompare, using the error propagated means and variances (on log10 scales) and Tukey’s HSD criterion. Where data was below the limit of detection, we tested for difference from that limit of detection.

## Supporting Information

Three supplementary figures (Figure S1 is a more detailed version of Figure 2a,b, Figure S2 depicts optimization of array assembly, and Figure S3 shows construction of a 4-spacer FnCas12a array), two supplementary tables (Table S1 includes oligos/primers used, and Table S2 includes CRISPR spacers used), and annotated DNA sequences for both the linear, genomically integrating vector pp2.1-CRISPR_8xkan_-Spec-pp2.2 and the replicating plasmid pBAV1spec-CRISPR_3xCRA_-Spec.

## Conflicts of Interest

R. Cooper and J. Hasty have filed a provisional patent 62/983,273 based on the technique described in this manusccript.

## Funding

Robert Cooper and Jeff Hasty were supported by the National Institutes of Health grant #R01-GM069811.

## For Table of Contents Use Only

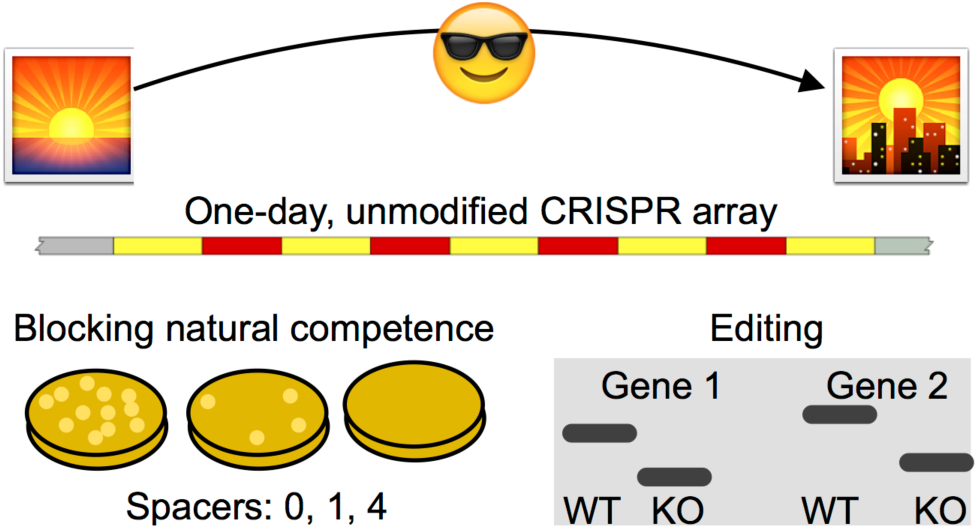

