## Supplementary Information for "One-Day Construction Of Multiplex Arrays to Harness Natural CRISPR Systems"

### Supporting Information

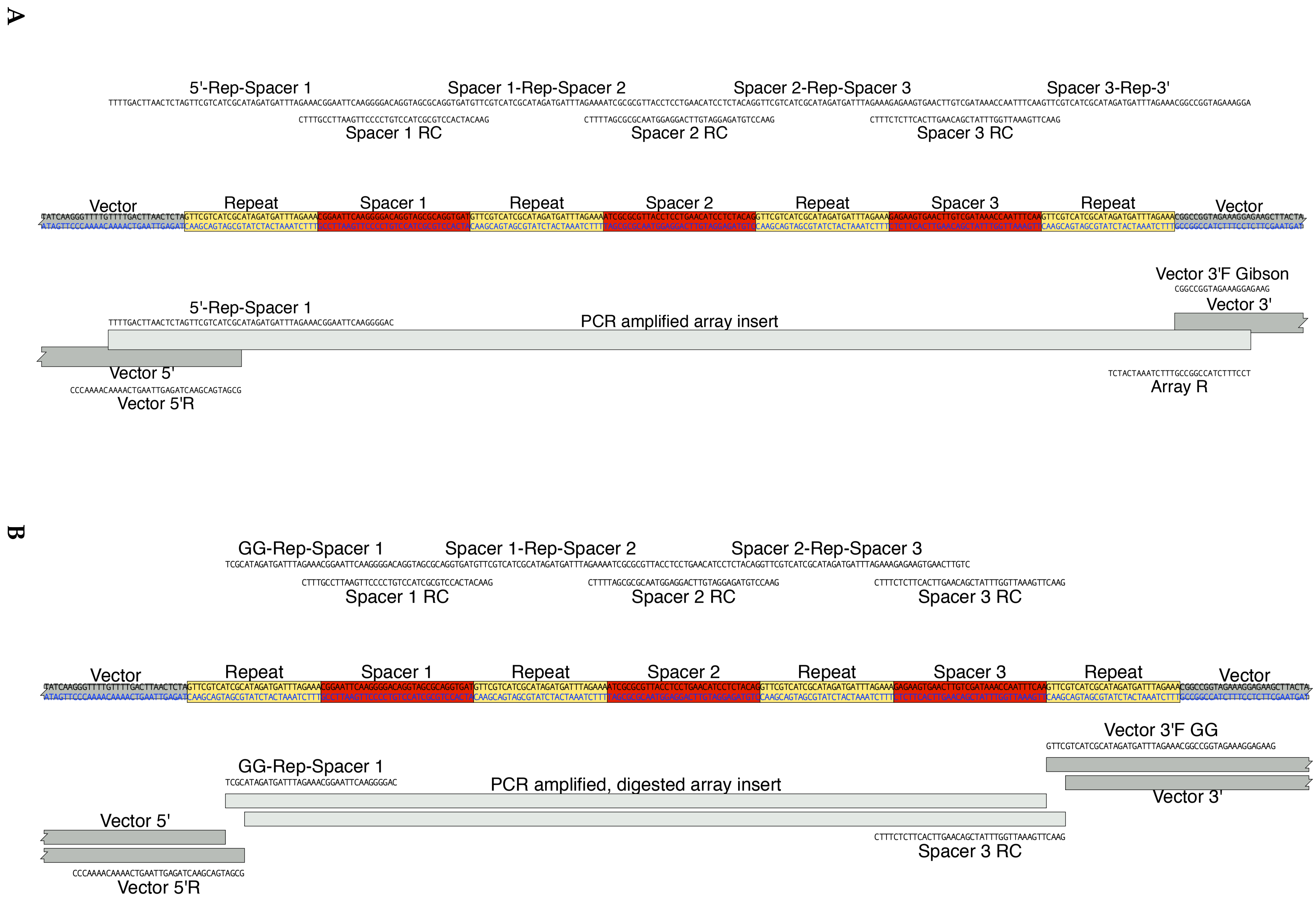
A
5'-Rep-Spacer 1
Spacer 1-Rep-Spacer 2
Spacer 2-Rep-Spacer 3
Spacer 3-Rep-3'
Spacer 1 RC
Spacer 2 RC
Spacer 3 RC
Vector
Repeat
Spacer 1
Repeat
Spacer 2
Repeat
Spacer 3
Repeat
Vector
Vector 3'F Gibson
Vector 3'
5'-Rep-Spacer 1
PCR amplified array insert
Vector 5'
Vector 5'R
Array R
B
GG-Rep-Spacer 1
Spacer 1-Rep-Spacer 2
Spacer 2-Rep-Spacer 3
Spacer 1 RC
Spacer 2 RC
Spacer 3 RC
Vector
Repeat
Spacer 1
Repeat
Spacer 2
Repeat
Spacer 3
Repeat
Vector
Vector 3'F GG
GG-Rep-Spacer 1
PCR amplified, digested array insert
Vector 3'
Vector 5'
Spacer 3 RC
Vector 5'R

**Figure S1: Detailed multiplex natural CRISPR array assembly.** A more detailed version of Figure 2 showing DNA sequences for the 3xBAP CRISPR array. The two array assembly strategies are for insertion into a vector using Gibson assembly or fusion PCR (A) or Golden Gate assembly (B). Note that primers used for Golden Gate assembly (denoted “GG” in B) have an additional BsaI site-containing tail appended to their 5′ ends that is not shown, specifically, TTTGGTCTCA.

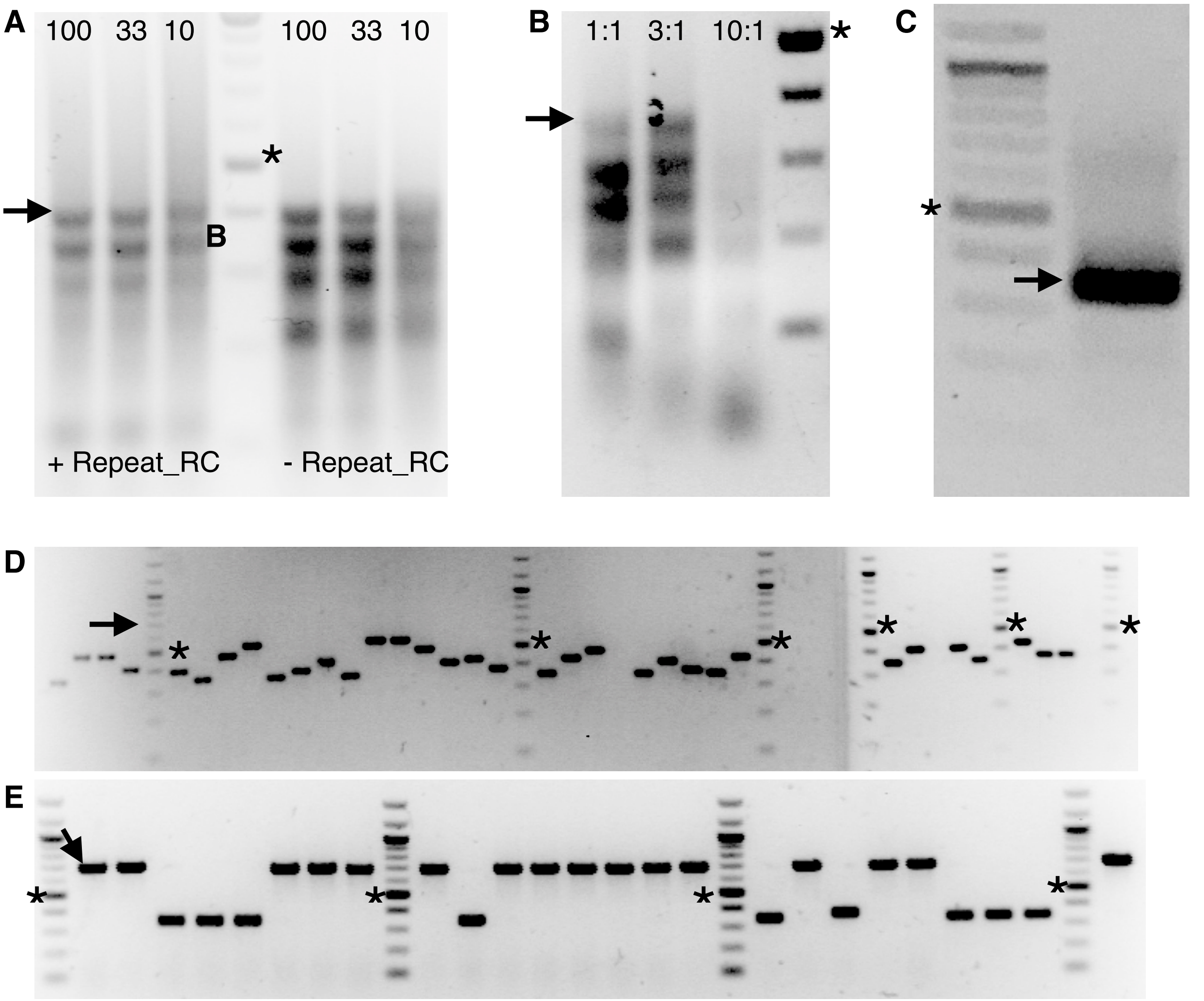


**Figure S2: Multiplex array assembly optimization.** Protocol optimizations were performed using a 6xIS-CRA array and inserted into pBAV using Golden Gate assembly. A,B: raw ligations, C: PCR amplification, D,E: Colony PCR screening of clones. Asterisks on all gels indicate the 500 bp band of the ladder, and arrows indicate the correctly sized assembly. A) Including the Repeat_RC oligo increases incorrect, higher-molecular-weight smearing (left 3 vs right 3 lanes), and 100 uM stock oligos (lanes 1 and 5) work better than 33 uM (lanes 2 and 6) or 10 uM (lanes 3 and 7) stock solutions. The center (lane 4) is a 100 bp ladder. B) Annealing and ligation is most efficient using 3 parts bottom oligos to 1 part top oligos. The lanes from left to right are ligations using 1:1, 3:1, and 10:1 ratios of bottom oligos to top oligos, followed by a 100 bp ladder. C) PCR amplification of the resulting ligation improves yield of the correct-sized product. D) Golden Gate assembly directly from ligation products yielded no correct-sized arrays out of 36 tested clones. All of 6 sequenced clones were correct at the 3′ end, but truncated at the 5′ end of the array. E) As for D, but the ligation product was PCR amplified and gel extracted before inserting into the vector. 16 of 25 colonies were the correct size, and all incorrect clones had 0x arrays (a single repeat only).

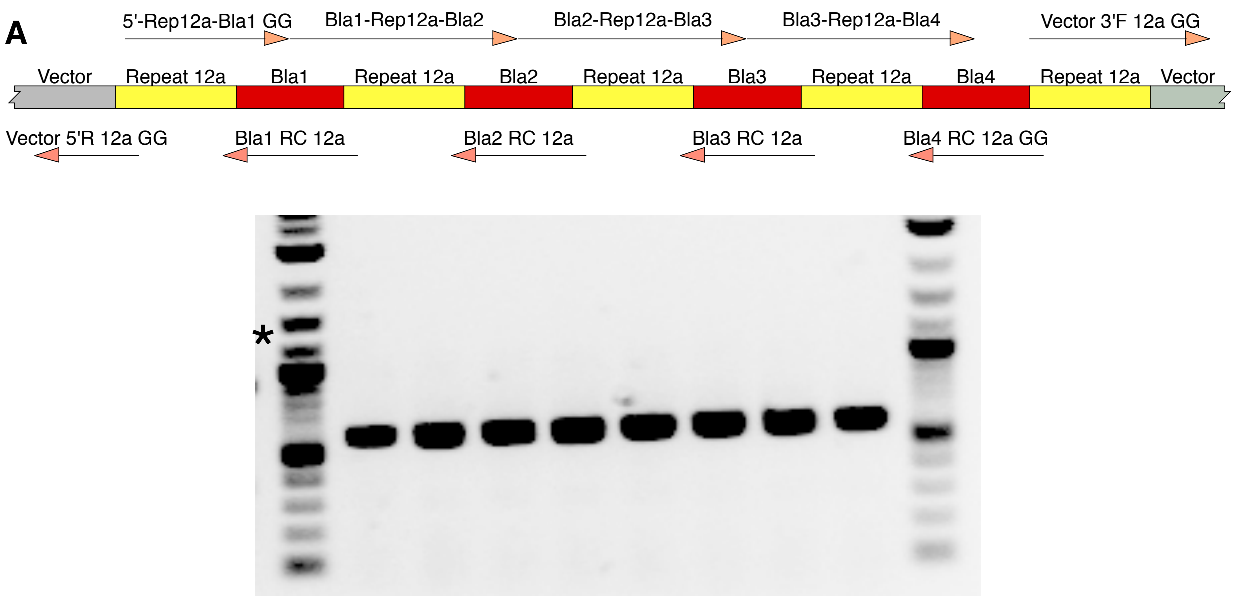


**Figure S3: Assembly of a 4-spacer Cas12a array.** A) Design and oligonucleotides for a 4-spacer FnCas12a CRISPR array, to be inserted into the vector using Golden Gate assembly. This is analogous to Figure 2B for *A. baylyi* arrays. All oligos denoted by GG also contain a 5′ Golden Gate tail (see Methods and Table S1). B) Screening of 8 clones for the 4-spacer array, of which all had the desired 603 bp product. The ladder on the end lanes contains 100 bp increments up to 1 kb, which is marked with an asterisk.

**Table S1: Oligos/Primers**

| Purpose | Name | Sequence |
|---|---|---|
| CRISPR$_{4xKan1}$ Gibson | kan1 5'-R-T1 | TTTTGACTTAACTCTAGTTCGTCATCGCATAGATGATTTAGAA AGGTCGATCAGGGAGGA |
| | kan1 T1-R-B1 | TATCGGGGAAGAACAGGTTCGTCATCGCATAGATGATTTAGA AATTGCATTCTAAAACCT |
| | kan1 B1-R-T2 | TAAATACAGAAAACAGGTTCGTCATCGCATAGATGATTTAGAA AGTCGATACTATGTTAT |
| | kan1 T2-R-B2 | ACGCCAACTTTGAAAAGTTCGTCATCGCATAGATGATTTAGA AAAAGCGAGCTCGGTACT |
| | kan1 B2-R-3' | AAAACAATTCATCCAGGTTCGTCATCGCATAGATGATTTAGAA ACGGCCGGTAGAAAGGA |
| | Kan1 T1 RC | GAACCTGTTCTTCCCCGATATCCTCCCTGATCGACCTTTC |
| | Kan1 B1 RC | GAACCTGTTTTCTGTATTTAAGGTTTTAGAATGCAATTTC |
| | Kan1 T2 RC | GAACTTTTCAAAGTTGGCGTATAACATAGTATCGACTTTC |
| | Kan1 B2 RC | GAACCTGGATGAATTGTTTTAGTACCGAGCTCGCTTTTTC |
| CRISPR$_{4xKan2}$ Gibson | kan2 5'-R-T1 | TTTTGACTTAACTCTAGTTCGTCATCGCATAGATGATTTAGAA ATCGCCGTCGGGCATGC |
| | kan2 T1-R-B1 | GCGCCTTGAGCCTGGCGTTCGTCATCGCATAGATGATTTAGA AAGGCTACCTGCCCATTC |
| | kan2 B1-R-T2 | GACCACCAAGCGAAACGTTCGTCATCGCATAGATGATTTAGA AACAACCTTACCAGAGGG |
| | kan2 T2-R-B2 | CGCCCCAGCTGGCAATGTTCGTCATCGCATAGATGATTTAGA AAGGCCGCTTGGGTGGAG |
| | kan2 B2-R-3' | AGGCTATTCGGCTATGGTTCGTCATCGCATAGATGATTTAGAA ACGGCCGGTAGAAAGGA |
| | Kan2 T1 RC | GAACGCCAGGCTCAAGGCGCGCATGCCCGACGGCGATTTC |
| | Kan2 B1 RC | GAACGTTTCGCTTGGTGGTCGAATGGGCAGGTAGCCTTTC |
| | Kan2 T2 RC | GAACATTGCCAGCTGGGGCGCCCTCTGGTAAGGTTGTTTC |
| | Kan2 B2 RC | GAACCATAGCCGAATAGCCTCTCCACCCAAGCGGCCTTTC |
| Array PCR | Array R | TCCTTTCTACCGGCCGTTTCTAAATCATCT |
| | Array R GG | TTTGGTCTCATCCTTTCTACCGGCCGTTTCTAAATCATCT |
| CRISPR$_{8xKan}$ Gibson | Kan1 B2-R-Kan2 T1 | AAAACAATTCATCCAGGTTCGTCATCGCATAGATGATTTAGAA ATCGCCGTCGGGCATGC |
| Vector Gibson | Vector 5'R | TAGAGTTAAGTCAAAACAAAACCC |

| | | |
|---|---|---|
| | Vector 3'F | GAAACGGCCGGTAGAAAGGA |
| Vector Golden Gate (GG) | Vector 5'R GG | TTTGGTCTCAGCGATGACGAACTAGAGTTAAGTCAAAACAAAACCC |
| | Vector 3'F GG | TTTGGTCTCAGTTCGTCATCGCATAGATGATTTAGAAACGGCCGGTAGAAAGGAGAAG |
| Genomic integrating CRISPR vector | pp 5'F | TGAGCCGACATTTTATTACCCTCT |
| | pp 3'R | TTACCTGAAAGCCAATCGCTG |
| CRISPR$_{3xBAP}$ GG | GG-R-BAP1 | TTTGGTCTCATCGCATAGATGATTTAGAAACGGAATTCAAGGGGAC |
| | BAP1-R-BAP2 | AGGTAGCGCAGGTGATGTTCGTCATCGCATAGATGATTTAGAAAATCGCGCGTTACCTCC |
| | BAP2-R-BAP3 | TGAACATCCTCTACAGGTTCGTCATCGCATAGATGATTTAGAAAGAGAAGTGAACTTGTC |
| | BAP1 RC | GAACATCACCTGCGCTACCTGTCCCCTTGAATTCCGTTTC |
| | BAP2 RC | GAACCTGTAGAGGATGTTCAGGAGGTAACGCGCGATTTTC |
| | BAP3 RC GG | TTTGGTCTCAGAACTTGAAATTGGTTTATCGACAAGTTCACTTCTCTTTC |
| CRISPR$_{3xCRA\text{-}3xBAP}$ GG | GG-R-CRA1 | TTTGGTCTCATCGCATAGATGATTTAGAAATCTCCGCGCTTGCTTC |
| | CRA1-R-CRA2 | GCATAATGCAGATTGAGTTCGTCATCGCATAGATGATTTAGAAAGTCACTATGACCATGT |
| | CRA2-R-CRA3 | TGCTTTGTATTGTGAAGTTCGTCATCGCATAGATGATTTAGAAACCCGGATTTTGACTGG |
| | CRA3-R-BAP1 | CGAAATGTAGAAGATAGTTCGTCATCGCATAGATGATTTAGAAACGGAATTCAAGGGGAC |
| | CRA1 RC | GAACTCAATCTGCATTATGCGAAGCAAGCGCGGAGATTTC |
| | CRA2 RC | GAACTTCACAATACAAAGCAACATGGTCATAGTGACTTTC |
| | CRA3 RC | GAACTATCTTCTACATTTCGCCAGTCAAAATCCGGGTTTC |
| PCR screening of arrays | Array screen F | GGAGTTCTGAGGTCATTACTGGATCTA |
| | Array screen R | CAAATGTACGGCCAGCAACG |
| *bap* deletion donor DNA | BAP 5'F | AGCAGCTGAGAGCCTGAATG |
| | BAP 5'R | ACATGCCAGCACTTAATCTGA |

| | | |
|---|---|---|
| | BAP 3’F | TCAGATTAAGTGCTGGCATGTGCACCCAATCCCTAACATTAAACA |
| | BAP 3'R | GGTTCGGGCACCTCATCATT |
| CRAΦ deletion donor DNA | CRA 5’F | ACAGGGCAGCCATTAACTGA |
| | CRA 5'R | TCTGAGACTGTAGCCTACGCA |
| | CRA 3’F | TGCGTAGGCTACAGTCTCAGAACGAAGTTATGTGCCACAAGAAA |
| | CRA 3'R | TCAGACGCAAGCGTGAAGAT |
| *bap* deletion screening | BAP checkF | GCCTCCTAAAATTGGGGGCT |
| | BAP checkR | CTTGGTTCTGCATTGGGTGC |
| CRAΦ deletion screening | CRA checkF | GACTTGCGTAGGCTTGGACT |
| | CRA checkR | GCATGTCATGGTTTGGTGGG |
| | CRA circular F | ATGAACGCGATCATTGCAGC |
| | CRA circular R | TACGGCCAATTGATCACCCA |
| Cas12a/Cpf1 $CRISPR_{4xBla}$ array | GG-R12a-Bla1 | TTTGGTCTCATAAGAACTTTAAATAATTTCTACTGTTGTAGATCGGCGTCAATACGGGA |
| | Bla1-R12a-Bla2 | TAATACCGCGCCACATGTCTAAGAACTTTAAATAATTTCTACTGTTGTAGATGGAGCTGAATGAAGCC |
| | Bla2-R12a-Bla3 | ATACCAAACGACGAGCGTCTAAGAACTTTAAATAATTTCTACTGTTGTAGATCTCCCGTATCGTAGTT |
| | Bla3-R12a-Bla4 | ATCTACACGACGGGGAGTCTAAGAACTTTAAATAATTTCTACTGTTGTAGATAGCCGGAAGGGCCGAG |
| | Vector 3’F 12a GG | TTTGGTCTCAGTCTAAGAACTTTAAATAATTTCTACTGTTGTAGATCGGCCGGTAGAAAGGACA |
| | Vector 5'R 12a GG | TTTGGTCTCACTTAGACTAGAGTTAAGTCAAAACAAAACCC |
| | Bla1 RC 12a | AGACATGTGGCGCGGTATTATCCCGTATTGACGCCGATCT |
| | Bla2 RC 12a | AGACGCTCGTCGTTTGGTATGGCTTCATTCAGCTCCATCT |
| | Bla3 RC 12a | AGACTCCCCGTCGTGTAGATAACTACGATACGGGAGATCT |
| | Bla4 RC 12a | TTTGGTCTCAAGACCAGGACCACTTCTGCGCTCGGCCCTTCCGGCTATCT |

#### Table S2: CRISPR spacers

| Name | Sequence |
|---|---|
| Kan1_T1 | GGTCGATCAGGGAGGATATCGGGGAAGAACAG |
| Kan1_T2 | GTCGATACTATGTTATACGCCAACTTTGAAAA |
| Kan1_B1 | TTGCATTCTAAAACCTTAAATACAGAAAACAG |
| Kan1_B2 | AAGCGAGCTCGGTACTAAAACAATTCATCCAG |
| Kan2_T1 | TCGCCGTCGGGCATGCGCGCCTTGAGCCTGGC |
| Kan2_T2 | CAACCTTACCAGAGGGCGCCCCAGCTGGCAAT |
| Kan2_B1 | GGCTACCTGCCCATTCGACCACCAAGCGAAAC |
| Kan2_B2 | GGCCGCTTGGGTGGAGAGGCTATTCGGCTATG |
| CRA1 | TCTCCGCGCTTGCTTCGCATAATGCAGATTGA |
| CRA2 | GTCACTATGACCATGTTGCTTTGTATTGTGAA |
| CRA3 | CCCGGATTTTGACTGGCGAAATGTAGAAGATA |
| BAP1 | CGGAATTCAAGGGGACAGGTAGCGCAGGTGAT |
| BAP2 | ATCGCGCGTTACCTCCTGAACATCCTCTACAG |
| BAP3 | GAGAAGTGAACTTGTCGATAAACCAATTTCAA |
| Random | TAGGGGAAAGCCTACTAGCCGGAGTGTTGCGA |

#### DNA sequence of sample genomically integrating vector, pp2.1-CRISPR$_{8xKan}$-Spec-pp2.2

LOCUS pp2.1-CR_4xAPH4x 3872 bp ss-DNA linear SYN 03-Jun-2016
DEFINITION -
ACCESSION -
KEYWORDS -
SOURCE -
FEATURES Location/Qualifiers
misc_feature <1..1006
/note="ADP1 prophage 2.1 region 2,158,257-2,159,574
[Split]"
misc_feature 10007..1087
/note="ADP1 CRISPR upstream region"
primer_bind complement(1064..1099)
/note="CRISPR 5'R 65"
primer_bind 1072..1131
/note="APH 5'-R-ST1"
repeat_region 1088..1115
/note="CR Repeat"
primer_bind complement(1112..1151)
/note="APH ST1 RC"

```
primer_bind     1132..1191
                /note="APH ST1-R-SB1"
repeat_region   1148..1175
                /note="CR Repeat"
primer_bind     complement(1172..1211)
                /note="APH SB1 RC"
primer_bind     1192..1251
                /note="APH SB1-R-ST2"
repeat_region   1208..1235
                /note="CR Repeat"
primer_bind     complement(1212..1231)
                /note="Repeat RC"
primer_bind     complement(1232..1271)
                /note="APH ST2 RC"
primer_bind     1252..1311
                /note="APH ST2-R-SB2"
repeat_region   1268..1295
                /note="CR Repeat"
primer_bind     complement(1292..1331)
                /note="APH SB2 RC"
primer_bind     1312..1371
                /note="APHB2-R-RCKT1"
repeat_region   1328..1355
                /note="CR Repeat"
primer_bind     complement(1352..1391)
                /note="RCK T1 RC"
primer_bind     1372..1431
                /note="RCK T1-R-B1"
repeat_region   1388..1415
                /note="CR Repeat"
primer_bind     complement(1412..1451)
                /note="RCK B1 RC"
primer_bind     1432..1491
                /note="RCK B1-R-T2"
repeat_region   1448..1475
                /note="CR Repeat"
primer_bind     complement(1472..1511)
                /note="RCK T2 RC"
primer_bind     1492..1551
                /note="RCK T2-R-B2"
repeat_region   1508..1535
                /note="CR Repeat"
primer_bind     complement(1532..1571)
                /note="RCK B2 RC"
misc_feature    1536..1567
                /note="RCK B2"
primer_bind     1552..1611
                /note="RCK B2-R-spec"
repeat_region   1568..1595
                /note="CR Repeat"
primer_bind     1592..1611
                /note="Vector 3'F"
primer_bind     complement(1592..1611)
                /note="Array R"
promoter        1656..1684
                /note="PampR"
CDS             1719..2510
                /codon_start=1
```

```
                /db_xref="GI:336359759"
                /gene="specR"
                /note="spectinomycin resistance marker"
                /product="SpecR"
                /protein_id="AEI53620.1"
                /transl_table=11
                /translation="MREAVIAEVSTQLSEVVGVIERHLEPTLLAVHLYGSAVDGGLKPH
                SDIDLLVTVTVRLDETTRRALINDLLETSASPGESEILRAVEVTIVVHDDIIPWRYPAK
                RELQFGEWQRNDILAGIFEPATIDIDLAILLTKAREHSVALVGPAAEELFDPVPEQDLF
                EALNETLTLWNSPPDWAGDERNVVLTLSRIWYSAVTGKIAPKDVAADWAMERLPAQYQP
                VILEARQAYLGQEEDRLASRADQLEEFVHYVKGEITKVVGK"
     misc_feature    2848..3872
                /note="ADP1 prophage 2.2 region 2,161,721-2,162,745"
     primer_bind     complement(3852..3872)
                /note="pp2.2 R63"
BASE COUNT       1003 A       873 C       886 G      1110 T          0 OTHER
ORIGIN      ?
        1 TGAGCCGACA TTTTATTACC CTCTTATCAA ACCGTACCTT TCACATAACG AATGAATGAA
       61 TACCGTACAT GGAGTGCGGC CAACCCACAG CGAACATCAT ATTTCGCATC CATCACCGTA
      121 CGGTTTTCCG TTTTAAGCTC TGCCCATGAT CTATCATGGA AATAACGGCT AATGATCACC
      181 TGCATCCACT CAAGTGTCGT TTCACTGTCT GTACCATTAA TAATATCCAG TACTAAACGT
      241 TGTACGGCAC GAGCTTCATT ATCGTTAATC TGACACGACA CTTTGTGACG TATAGCTTGT
      301 TGTACTCCTT GAGCATCACA AAGGTAATAA GCAATAAGTT TAGCTCGATC TTTCTTCTTT
      361 ACACGTACCT TGGCTTTCTT CATTGCAATA GCAATCGGGC TATCGGAATA CTGTCCACCA
      421 CGACAAGAAC GTTGCCACGC TCCACATTGG CGTAACCAAT CTGGCAAATC ATACTTGCTC
      481 CAATCCACCG TCTGCATAAT GTGCACTGCT GTATTCATCT CATCACCTAA TTTGTTTCAA
      541 GTTAAATTTT ATAAGCGTTA TTGTTTTATG GTTCTGCCTG CTCCTCTACC GATCTAAAAC
      601 GACAAGTTTC GAGATAATCC AGTACTCGAA CTGCACCGCG TTTACCGTGT CGGTTTTTCA
      661 CTACAATCAG CTCTGTGATT CCCATCGGTT TGGTTGAGTC TTCTGGATCG GTTAGGGGGT
      721 TAACAAGGAT GATCTGGTCT GCATCTTGCT CGATTTGTCC AGATTCTTTG ATATCTGATG
      781 CTTTAGGACG TTTGCCCTTC TCTGCCTCAC GGTTGAGCTG TACCAGTGCA ATGACAGGAC
      841 ATTCAAACTC TTTCGCCATG GATTTTAATT CACGGCTGAT GGAACTGACT TCCTGAAAGC
      901 GATCTTTCTT GCTCGGGTCT CTGAGCAGTT GTAAATAATC CACGATGATG CAGCCCAATT
      961 CCTTGTAACG GCGTTTGGCT CGACGTGCAT AGGAACGGAC CTCACTCAAG TGATTCATAA
     1021 CGAAGTATTT TTACTCATTA AAAGCTTATA TAATTGATAT CAAGGGTTTT GTTTTGACTT
     1081 AACTCTAGTT CGTCATCGCA TAGATGATTT AGAAAGGTCG ATCAGGGAGG ATATCGGGGA
     1141 AGAACAGGTT CGTCATCGCA TAGATGATTT AGAAATTGCA TTCTAAAACC TTAAATACAG
     1201 AAAACAGGTT CGTCATCGCA TAGATGATTT AGAAAGTCGA TACTATGTTA TACGCCAACT
     1261 TTGAAAAGTT CGTCATCGCA TAGATGATTT AGAAAAAGCG AGCTCGGTAC TAAAACAATT
     1321 CATCCAGGTT CGTCATCGCA TAGATGATTT AGAAATCGCC GTCGGGCATG CGCGCCTTGA
     1381 GCCTGGCGTT CGTCATCGCA TAGATGATTT AGAAAGGCTA CCTGCCCATT CGACCACCAA
     1441 GCGAAACGTT CGTCATCGCA TAGATGATTT AGAAACAACC TTACCAGAGG GCGCCCCAGC
     1501 TGGCAATGTT CGTCATCGCA TAGATGATTT AGAAAGGCCG CTTGGGTGGA GAGGCTATTC
     1561 GGCTATGGTT CGTCATCGCA TAGATGATTT AGAAACGGCC GGTAGAAAGG AGAAGCTTAC
     1621 TAGCTATTTG TTTATTTTTC TAAATACATT CAAATATGTA TCCGCTCATG AGACAATAAC
     1681 CCTGATAAAT GCTTCAATAA TATTGAAAAA GGAAGAGTAT GAGGGAAGCG GTGATCGCCG
     1741 AAGTATCGAC TCAACTATCA GAGGTAGTTG GCGTCATCGA GCGCCATCTC GAACCGACGT
     1801 TGCTGGCCGT ACATTTGTAC GGCTCCGCAG TGGATGGCGG CCTGAAGCCA CACAGTGATA
     1861 TTGATTTGCT GGTTACGGTG ACCGTAAGGC TTGATGAAAC AACGCGGCGA GCTTTGATCA
     1921 ACGACCTTTT GGAAACTTCG GCTTCCCCTG GAGAGAGCGA GATTCTCCGC GCTGTAGAAG
     1981 TCACCATTGT TGTGCACGAC GACATCATTC CGTGGCGTTA TCCAGCTAAG CGCGAACTGC
     2041 AATTTGGAGA ATGGCAGCGC AATGACATTC TTGCAGGTAT CTTCGAGCCA GCCACGATCG
     2101 ACATTGATCT GGCTATCTTG CTGACAAAAG CAAGAGAACA TAGCGTTGCC TTGGTAGGTC
     2161 CAGCGGCGGA GGAACTCTTT GATCCGGTTC CTGAACAGGA TCTATTTGAG GCGCTAAATG
     2221 AAACCTTAAC GCTATGGAAC TCGCCGCCCG ACTGGGCTGG CGATGAGCGA AATGTAGTGC
     2281 TTACGTTGTC CCGCATTTGG TACAGCGCAG TAACCGGCAA AATCGCGCCG AAGGATGTCG
     2341 CTGCCGACTG GGCAATGGAG CGCCTGCCGG CCCAGTATCA GCCCGTCATA CTTGAAGCTA
     2401 GACAGGCTTA TCTTGGACAA GAAGAAGATC GCTTGGCCTC GCGCGCAGAT CAGTTGGAAG
```

```
    2461 AATTTGTCCA CTACGTGAAA GGCGAGATCA CCAAGGTAGT CGGCAAATAA TGTCTAACAA
    2521 TTCGTTCAAG CCGAGGGGCC GCAAGATCCG GCCACGATGA CCCGGTCGTC GGTTCAGGGC
    2581 AGGGTCGTTA AATAGCCGCT TATGTCTATT GCTGGTTTAC CGGTTTATTG ACTACCGGAA
    2641 GCAGTGTGAC CGTGTGCTTC TCAAATGCCT GAGGTTTCAG CAAAAAACCC CTCAAGACCC
    2701 GTTTAGAGGC CCCAAGGGGT TATGCTAGTT ATTGCTCAGC GGTGGCAGCA GCCTAGGTTA
    2761 ATTAAGCTGC GCTAGTAGAC GAGTCCATGT GCTGGCGTTC AAATTTCGCA GCAGCGGTTT
    2821 CTTTACCAGA CTCGACAAGC TTACTAGAGT GTTCATATTG ACCTCGCTTA GTGTGGTTAA
    2881 TACGCCGCTT CTTGTTACTG CAAGAGGCGG TTTTTTTATG GGTGTACACA TGACTGCACC
    2941 TGTTGATTCA GTTCAGATAG TGCTTTGTGC ATCTCATGGA TGACTCTGGC CATATCCAGT
    3001 GCTTCGCCTT GGGTAATTCG CCCATCGGCC ATCATTTCTT TAAACAGTGC TGATATATCG
    3061 CCCTTCTTGA TGCCTATGCA TAGGAAGGTA TCCATCAGAC TGGTATCTCG CTGGCTCTCG
    3121 GGTATGTCTG GCAGGTCAAT TGCCACCTTT CCGAGTCGTG CACACATTTC CTGCAATATC
    3181 CGATAGTCCC CTGTAATCTC CATCAGCTTG ACTGCCTCGA GCAATGTAAT GTGATGGGTA
    3241 TGTGTGTTTG GGTTGACCTT GCTATTGAGC ACCGCAGGGC TTTTGATGCC TAAACGTGAT
    3301 GCAAGTGCAG ATGCACCACC CAGAAAGTCG TGAACGGTGT GATAGGCAGC ATCTAATATG
    3361 TTCATGGCGG GTTCCTTTGA ACGTGTTTAT TAGATGGGTG CTGACATAAG ATTGGATTTA
    3421 TGGTTAGGAC GTAATTCAAT CCAAATATCT TCGTAATCAT CTGGAAATAA ATCTTTGCGA
    3481 GTGCATAGAC CTTTATCTTC AGCAATTACA GCTAAGCGGA TTTTGCGATC TCTGGGAATT
    3541 GCTTTCCATC CACTTACGGA TGCAGCGGTG ATACCTAAAA ATCTAGCAAC AGCAGTTACA
    3601 CCACCTAAAA GCTCAATAAA TTGATCATCA GTCATGTTGA TCTCCTAATT TTATTGCCTC
    3661 AATTATTAGG TATTCCTTAT ATTTATCAA TAGGAATACC TTATTTATTT TATGTTAGGA
    3721 TTTCCTAATA GACTAGGTAA GATCATGAAA ACATTAGCTG AACGACTTAA ATATGCGATG
    3781 GAAATTTTGC CACCTAAGAA AATCAAGGGT GTCGAACTTG CTCGTGTAGT TGGAGTTAAA
    3841 CCACCATCTG TCAGCGATTG GCTTTCAGGT AA
//
```

#### DNA sequence of sample replicating vector, pBAV1spec-CRISPR$_{3xCRA}$-Spec

```
LOCUS       pBAV1spec_CR_9xC        3162 bp    DNA     circular SYN 22-Mar-2018
DEFINITION  -
ACCESSION   -
KEYWORDS    -
SOURCE      -
FEATURES             Location/Qualifiers
     terminator      81..158
                     /note="t1"
     CDS             complement(194..892)
                     /codon_start=1
                     /db_xref="GI:336359729"
                     /gene="repA"
                     /note="replication initiator protein"
                     /product="RepA"
                     /protein_id="AEI53594.1"
                     /transl_table=11
                     /translation="MAIKNTKARNFGFLLYPDSIPNDWKEKLESLGVSMAVSPLHDMDE
                     KKDKDTWNSSDVIRNGKHYKKPHYHVIYIARNPVTIESVRNKIKRKLGNSSVAHVEILD
                     YIKGSYEYLTHESKDAIAKNKHIYDKKDILNINDFDIDRYITLDESQKRELKNLLLDIV
                     DDYNLVNTKDLMAFIRLRGAEFGILNTNDVKDIVSTNSSAFRLWFEGNYQCGYRASYAK
                     VLDAETGEIK"
     gene            complement(194..892)
                     /gene="repA"
     CDS             complement(959..1120)
                     /codon_start=1
                     /db_xref="GI:336359730"
                     /note="ORFC"
                     /product="hypothetical protein"
                     /protein_id="AEI53595.1"
```

/transl_table=11
/translation="MVISESKKRVMISLTKEQDKKLTDMAKQKGFSKSAVAALAIEEYA
RKESEQKK"
CDS complement(1161..1370)
/codon_start=1
/db_xref="GI:336359731"
/note="ORFB"
/product="hypothetical protein"
/protein_id="AEI53596.1"
/transl_table=11
/translation="MGGKEANFASVLRPPIKCRVPIFVPKTLYPNWLKGLRGFSIANES
PTFSPTFFINLYLSSFIVVFMITK"
repeat_region 1323..1371
/note="IRIII"
repeat_region 1455..1477
/note="IRII"
repeat_region 1524..1655
/note="IRI"
terminator 1708..>1799
/note="t0"
primer_bind 1773..1799
/note="Array screen F"
misc_feature 1801..1880
/note="CRISPR upstream reagion"
primer_bind complement(1857..1892)
/note="Vector R"
primer_bind 1865..1924
/note="5'-R-CRA1"
repeat_region 1881..1908
/note="CR Repeat"
primer_bind complement(1905..1944)
/note="CRA targ1 RC"
primer_bind 1925..1984
/note="CRA1-R-CRA2"
repeat_region 1941..1968
/note="CR Repeat"
primer_bind complement(1965..2004)
/note="CRA targ2 RC"
/note="CRA2-R-CRA3"
repeat_region 2001..2028
/note="CR Repeat"
primer_bind complement(2025..2064)
/note="CRA targ3 RC"
primer_bind 2045..>2088
/note="CRA3-R-PP1"
repeat_region 2061..2088
/note="CR Repeat"
primer_bind complement(2075..2104)
/note="Array R"
primer_bind 2085..2104
/note="Vector F"
promoter 2149..2177
/note="PampR"
CDS 2212..3003
/codon_start=1
/db_xref="GI:336359759"
/gene="specR"
/note="spectinomycin resistance marker"

/product="SpecR"
/protein_id="AEI53620.1"
/transl_table=11
/translation="MREAVIAEVSTQLSEVVGVIERHLEPTLLAVHLYGSAVDGGLKPH
SDIDLLVTVTVRLDETTRRALINDLLETSASPGESEILRAVEVTIVVHDDIIPWRYPAK
RELQFGEWQRNDILAGIFEPATIDIDLAILLTKAREHSVALVGPAAEELFDPVPEQDLF
EALNETLTLWNSPPDWAGDERNVVLTLSRIWYSAVTGKIAPKDVAADWAMERLPAQYQP
VILEARQAYLGQEEDRLASRADQLEEFVHYVKGEITKVVGK"
primer_bind complement(2291..2310)
/note="Array screen R"
BASE COUNT 899 A 688 C 627 G 948 T 0 OTHER
ORIGIN ?
1 TAGAAAGGAG AAGCTTACTA GTAGCGGCCG CTGCAGGCCT CAGGGCCCGA TCGATGCCGC
61 CGCTTAATTA ATTAATCCAG AGGCATCAAA TAAAACGAAA GGCTCAGTCG AAAGACTGGG
121 CCTTTCGTTT TATCTGTTGT TTGTCGGTGA ACGCTCTCCT GAGTAGGACA AATCCGCCGC
181 CCTAGACCTA GTGTCATTTT ATTTCCCCCG TTTCAGCATC AAGAACCTTT GCATAACTTG
241 CTCTATATCC ACACTGATAA TTGCCCTCAA ACCATAATCT AAAGGCGCTA GAGTTTGTTG
301 AAACAATATC TTTTACATCA TTCGTATTTA AAATTCCAAA CTCCGCTCCC CTAAGGCGAA
361 TAAAAGCCAT TAAATCTTTT GTATTTACCA AATTATAGTC ATCCACTATA TCTAAGAGTA
421 AATTCTTCAA TTCTCTTTTT TGGCTTTCAT CAAGTGTTAT ATAGCGGTCA ATATCAAAAT
481 CATTAATGTT CAAAATATCT TTTTTGTCGT ATATATGTTT ATTCTTAGCA ATAGCGTCCT
541 TTGATTCATG AGTCAAATAT TCATATGAAC CTTTGATATA ATCAAGTATC TCAACATGAG
601 CAACTGAACT ATTCCCCAAT TTTCGCTTAA TCTTGTTCCT AACGCTTTCT ATTGTTACAG
661 GATTTCGTGC AATATATATA ACGTGATAGT GTGGTTTTTT ATAGTGCTTT CCATTTCGTA
721 TAACATCACT ACTATTCCAT GTATCTTTAT CTTTTTTTTC GTCCATATCG TGTAAAGGAC
781 TGACAGCCAT AGATACGCCC AAACTCTCTA ATTTTTCCTT CCAATCATTA GGAATTGAGT
841 CAGGATATAA TAAAAATCCA AAATTTCTAG CTTTAGTATT TTTAATAGCC ATGATATAAT
901 TACCTTATCA AAAACAAGTA GCGAAAACTC GTATCCTTCT AAAAACGCGA GCTTTCGCTT
961 ATTTTTTTTG TTCTGATTCC TTTCTTGCAT ATTCTTCTAT AGCTAACGCC GCAACCGCAG
1021 ATTTTGAAAA ACCTTTTTGT TTCGCCATAT CTGTTAATTT TTTATCTTGC TCTTTTGTCA
1081 GAGAAATCAT AACTCTTTTT TTCGATTCTG AAATCACCAT TTAAAAAACT CCAATCAAAT
1141 AATTTTATAA AGTTAGTGTA TCACTTTGTA ATCATAAAAA CAACAATAAA GCTACTTAAA
1201 TATAGATTTA TAAAAAACGT TGGCGAAAAC GTTGGCGATT CGTTGGCGAT TGAAAAACCC
1261 CTTAAACCCT TGAGCCAGTT GGGATAGAGC GTTTTTGGCA CAAAAATTGG CACTCGGCAC
1321 TTAATGGGGG GTCGTAGTAC GGAAGCAAAA TTCGCTTCCT TTCCCCCCAT TTTTTTCCAA
1381 ATTCCAAATT TTTTTCAAAA ATTTTCCAGC GCTACCGCTC GGCAAAATTG CAAGCAATTT
1441 TTAAAATCAA ACCCATGAGG GAATTTCATT CCCTCATACT CCCTTGAGCC TCCTCCAACC
1501 GAAATAGAAG GGCGCTGCGC TTATTATTTC ATTCAGTCAT CGGCTTTCAT AATCTAACAG
1561 ACAACATCTT CGCTGCAAAG CCACGCTACG CTCAAGGGCT TTTACGCTAC GATAACGCCT
1621 GTTTTAACGA TTATGCCGAT AACTAAACGA AATAAACGCT AAAACGTCTC AGAAACGATT
1681 TTGAGACGTT TTAATAAAAA ATCGCCTAGT GCTTGGATTC TCACCAATAA AAAACGCCCG
1741 GCGGCAACCG AGCGTTCTGA ACAAATCCAG ATGGAGTTCT GAGGTCATTA CTGGATCTAC
1801 AAGTGATTCA TAACGAAGTA TTTTTACTCA TTAAAAGCTT ATATAATTGA TATCAAGGGT
1861 TTTGTTTTGA CTTAACTCTA GTTCGTCATC GCATAGATGA TTTAGAAATC TCCGCGCTTG
1921 CTTCGCATAA TGCAGATTGA GTTCGTCATC GCATAGATGA TTTAGAAAGT CACTATGACC
1981 ATGTTGCTTT GTATTGTGAA GTTCGTCATC GCATAGATGA TTTAGAAACC CGGATTTTGA
2041 CTGGCGAAAT GTAGAAGATA GTTCGTCATC GCATAGATGA TTTAGAAACG GCCGGTAGAA
2101 AGGAGAAGCT TACTAGCTAT TTGTTTATTT TTCTAAATAC ATTCAAATAT GTATCCGCTC
2161 ATGAGACAAT AACCCTGATA AATGCTTCAA TAATATTGAA AAAGGAAGAG TATGAGGGAA
2221 GCGGTGATCG CCGAAGTATC GACTCAACTA TCAGAGGTAG TTGGCGTCAT CGAGCGCCAT
2281 CTCGAACCGA CGTTGCTGGC CGTACATTTG TACGGCTCCG CAGTGGATGG CGGCCTGAAG
2341 CCACACAGTG ATATTGATTT GCTGGTTACG GTGACCGTAA GGCTTGATGA AACAACGCGG
2401 CGAGCTTTGA TCAACGACCT TTTGGAAACT TCGGCTTCCC CTGGAGAGAG CGAGATTCTC
2461 CGCGCTGTAG AAGTCACCAT TGTTGTGCAC GACGACATCA TTCCGTGGCG TTATCCAGCT
2521 AAGCGCGAAC TGCAATTTGG AGAATGGCAG CGCAATGACA TTCTTGCAGG TATCTTCGAG
2581 CCAGCCACGA TCGACATTGA TCTGGCTATC TTGCTGACAA AAGCAAGAGA ACATAGCGTT
2641 GCCTTGGTAG GTCCAGCGGC GGAGGAACTC TTTGATCCGG TTCCTGAACA GGATCTATTT
2701 GAGGCGCTAA ATGAAACCTT AACGCTATGG AACTCGCCGC CCGACTGGGC TGGCGATGAG

2761 CGAAATGTAG TGCTTACGTT GTCCCGCATT TGGTACAGCG CAGTAACCGG CAAAATCGCG
2821 CCGAAGGATG TCGCTGCCGA CTGGGCAATG GAGCGCCTGC CGGCCCAGTA TCAGCCCGTC
2881 ATACTTGAAG CTAGACAGGC TTATCTTGGA CAAGAAGAAG ATCGCTTGGC CTCGCGCGCA
2941 GATCAGTTGG AAGAATTTGT CCACTACGTG AAAGGCGAGA TCACCAAGGT AGTCGGCAAA
3001 TAATGTCTAA CAATTCGTTC AAGCCGAGGG GCCGCAAGAT CCGGCCACGA TGACCCGGTC
3061 GTCGGTTCAG GGCAGGGTCG TTAAATAGCC GCTTATGTCT ATTGCTGGTT TACCGGTTTA
3121 TTGACTACCG GAAGCAGTGT GACCGTGTGC TTCTCAAATG CC
//
